# *In silico* screening of TMPRSS2 SNPs that affect its binding with SARS-CoV2 spike protein and directly involved in the interaction affinity changes

**DOI:** 10.1101/2021.09.29.462283

**Authors:** Fatma Nouira, Manel Hamdi, Alaeddine Redissi, Soumaya Kouidhi, Cherine Charfeddine, Meriem M’saad, Ameur Cherif, Sabri Messaoudi, Sarah Aldulaijan, Noureddine Raouafi, Adnene Dhouib, Amor Mosbah

## Abstract

In this paper, we used in silico analysis to shed light on the possible interaction between TMPRSS2 and SARS-CoV2 spike (S) protein by examining the role of TMPRSS2 single nucleotide polymorphisms (SNPs) in relation with susceptibility and inter-individual variability of SARS-CoV2 infection. First, we used molecular docking of human TMPRSS2 protein to predict the binding site of TMPRSS2, especially the TMPRSS2 link loops, in order to assess the effect TMPRSS2 SNPs. The latter lead to missense variants on the interaction between TMPRSS2 and SARS-CoV2 S protein. In a second step, we further refine our analysis by performing a structure-function analysis of the complexes using PyMol software, and finally by MD simulations to validate the as-obtained results. Our findings show that 17 SNPs among the 692 natural TMPRSS2 coding variants are in positions to influence the binding of TMPRSS2 with the viral S protein. All of them give more important interaction energy as assessed by docking. Among the 17 SNPs, four missense variants E389A, K392Q, T393S and Q438E lead to “directly increasing” the interaction affinity and 2 missense variants R470I and Y416C cause it “directly decreasing”. The R470I and Y416C present in African and American population, respectively. While the other 4 SNP variants (E389A; K392Q; T393S and Q438E) are present only in the European population, which could link the viral infection susceptibility to demographic, geographic and genetic factors.

## INTRODUCTION

The pathogenesis of the coronavirus disease (COVID-19) is triggered by the entry of SARS-CoV2 via the spike protein into angiotensin-converting enzyme 2 (ACE2)-bearing host cells, especially pneumocytes, resulting in overactivation of the immune system (cytokine storm), which attacks the infected cells and damages the lung tissue (Hakmi et al.,2020). Cell entry of the betacoronaviruses, depends on the binding of the surface unit, S1, of the viral spike protein to ACE2 receptor, which facilitates viral attachment to the surface of the target cells. Moreover, to fuse membranes, the S protein needs to be proteolytically activated at the S1/S2 boundary, such that S1 dissociates and S2 undergoes a radical structural modification, therefore, viral entry requires not only the binding to the ACE2 receptor but also the priming of the viral S protein by the transmembrane protease serine 2 (TMPRSS2), which cleaves the S proteins at the S1/S2 and S2 sites (Hoffman et al., 2020; Baughn et al., 2020). This step is mandatory for the virus-host cell membrane fusion and cell entry (Hoffman et al., 2020; Matsuyama et al., 2020).

TMPRSS2 is an essential enzyme that can cleave hemagglutinin of many subtypes of the influenza virus and the coronavirus S protein including severe acute respiratory syndrome-related coronavirus (SARS-CoV) (Hoffmann et al., 2020) and the Middle East respiratory syndrome-related coronavirus (MERS-CoV) (Du et al., 2017), thus facilitating the virus-cell membrane fusion and viral infection (Böttcher et al., 2006). Matsuyama *et al*. demonstrated that TMPRSS2-expressing cell lines are highly susceptible to SARS-CoV, MERS-CoV, and SARS-CoV2 (Matsuyama et al., 2020), which proves that *TMPRSS2* expression is crucial for the spread of the virus and pathogenesis. Results from several studies on prostate cancer revealed that overexpression of *TMPRSS2* induced by transactivation of androgen receptor caused growth, invasion and metastasis of prostate cancer stem cells (Chen et al., 2019; Ko et al., 2015). Recently, a plethora of evidence showed that single nucleotide polymorphisms (SNPs) in *TMPRSS2* gene may be involved in several disorders including prostate and breast cancers via modulation of *TMPRSS2* expression (Bhanushali et al., 2018; Luostari et al., 2014). As a result, genetic variation in this gene may modulate genetic predisposition to infection and virus clearance in the host.

Most recently, the ongoing COVID-19 pandemic has created the hypothesis that inter-individual genetic differences may affect the spatial transmission dynamics of SARS-CoV2, the susceptibility and severity of disease, and the inflammatory and immune response (Paniri et al., 2020). Specifically, there is evidence that the TMPRSS2 plays a crucial role in SARS-CoV2 infection and it was speculated that *TMPRSS2* gene polymorphism may modulate the interaction between TMPRSS2 and the virus spike protein during the virus entry into the host cell and may influence individual’s susceptibility to the virus infection (Paniri et al., 2020; David et al., 2020; Singh et al., 2020). Furthermore, recent studies used statistical analysis and *in silico* tools to predict possible impact of an amino acid substitution/deletion on the structure and function of a given human protein to identify variants that could result in TMPRSS2 loss of structure/function and suggested that these variants may indirectly modulate the interaction affinity between TMPRSS2 and the invading virus (Zarubin et al., 2020; Vashnubhotla et al., 2020).

In this study, we used several bioinformatics tools and databases for a computational analysis of TMPRSS2 to determine the role of single nucleotide polymorphisms in susceptibility and inter-individual variability of SARS CoV2 infection by examining the effect of TMPRSS2 SNPs on the interaction of this protein with the S1/S2 domain of the spike protein. However, the molecular structure of human TMPRSS2 protein is not available in the protein database (PDB) and structural details of intermolecular interactions between SARS-CoV2 and TMPRSS2 are not very clear. So, we built the TMPRSS2 3D structure using I-TASSER, we predicted the binding site of TMPRSS2 protein, more specifically, all TMPRSS2 link loops and we have used *in silico* molecular docking to analyze the possible effects of TMPRSS2 SNPs leading to missense variants on the interaction between TMPRSS2 and the viral S protein. To further refine our analysis, we performed a structure function analysis of the complexes obtained by molecular docking using PyMol software (DeLano et al., 2002), followed by MD simulations using NAMD2 and VMD visualization software (Humphrey et al., 1996; Phillips et al., 2005) to validate the results obtained. To sum up, the idea of this approach is to detect the TMPRSS2 polymorphisms affecting binding interfaces, and which are directly associated with the increase or decrease of the interaction affinity with the S1/S2 domain of the spike protein, which can be considered as protective or susceptibility variants to SARS CoV-2 infection.

## MATERIAL AND METHODS

### TMPRSS2 polymorphism analysis

SNPs in *TMPRSS2*, with minor allele frequency (MAF) between 0.01 and 0.5, were extracted from Ensembl genome browser (https://asia.ensembl.org/index.html) (Cunningham et al., 2019), gnomAD (https://gnomad.broadinstitute.org/) (Karczewski et al, 2020), 1000 Genomes (https://www.internationalgenome.org/1000-genomes-browsers/) (Siva, 2008), and NHLBI (https://evs.gs.washington.edu/EVS/) (Auer et al., 2012) databases. Appropriate filters were employed to limit the data to only the missense and damaging SNPs of *TMPRSS2*. The functional impact of allelic variants of TMPRSS2 was predicted using sorting intolerant from tolerant (SIFT) (https://sift.bii.a-star.edu.sg/), which predicts the effects of amino acids substitution on protein structure, the score ranges of 0 to 0.05 are considered as deleterious substitutions (Ng et al., 2003), PolyPhen-2(http://genetics.bwh.harvard.edu/pph2/), is a useful database that predicts the possible consequences of amino acid substitution on functional and structural proteins. Score range of 0.0 – 0.15, 0.15 – 0.85 and 0.85 – 1.0 are considered benign, possibly damaging and damaging, respectively (Adzhubei et al., 2013). Combined annotation-dependent depletion (CADD) (https://cadd.gs.washington.edu) is a tool used to assess the harmfulness of single nucleotide variants in the human genome. Variants with CADD scores > 20 are considered deleterious variants (Rentzsch et al., 2019).

### Protein molecular modelling

When this study was started, the crystal structure of human TMPRSS2 has not been filed in the Protein Data Bank (PDB), therefore, we modelled the structure of human TMPRSS2 employing I-TASSER (Iterative Threading Assembly Refinement), which is a strong predictor of protein 3D structure, aiming to determine by computational calculations the spatial location of every atom in a protein molecule from the amino acid sequence (Zhang, 2008). In April 2021, the crystal structure of human TMPRSS2 in complex with Nefamostat has been deposited in the Protein Data Bank (code PDB: 7MEQ), we compared our structure to the one recently deposited in PDB in order to verify the quality and reliability of our model using PyMOL software.

### Identification of TMPRSS2 binding interfaces, selection and characterization of SNPs

Although there is not enough information about the active site and the catalytic site of TMPRSS2, by running a protease conserved domain (CD), TMPRSS2 was analyzed, and all its link loops residues were predicted with PyMOL. Following the identification of the binding interfaces, we selected only the variants located at the level of these connecting loops from the list of missense and damaging TMPRSS2 SNPs already predicted and extracted from databases. dbSNP is a database of genetic variants implemented at the National Center for Biotechnology Information “NCBI” and GnomaAD database were exploited to characterize the selected SNPs (population and allelic frequency).

### Homology modelling of selected TMPRSS2 SNPs affecting binding interfaces

To explore the structural changes in the protein encoded by different alleles of TMPRSS2, molecular models of all the selected protein variants were developed and superimposed over the structurally resolved template of wild-type TMPRSS2 using SWISS-MODEL, which allows a fully automated protein structure homology modelling. The FASTA sequence of TMPRSS2 was obtained from the UniProt protein knowledge database (UniProt Id O15393, corresponding to 492 amino acid transcript). The sequence of each TMPRSS2 variant is generated at the base of the wild-type sequence by a simple substitution of the amino acid coding for the missense mutation.

### Molecular docking

AutoDock Vina was used to carry out the molecular docking between S1/S2 domain of SARS-CoV2 **s**pike protein and TMPRSS2 wild type or missense variants. In our analysis we used, as a receptor, the TMPRSS2 wild type or missense variants, and, as a ligand, the S1/S2 domain of SARS-CoV2 spike protein model (Code PDB :6ZB4) downloaded from RCSB-PDB database. To obtain the optimal docking, the interactions of the wild-type receptor and variants with the partner were simulated using different parameters therefore receptor and ligand we used a grid size was set to 80× 80 × 80 points with a spacing of 1 Å.

### Structure analysis of TMPRSS2 variants and SARS-CoV2 spike protein complexes

To further understand the effect of polymorphisms on receptor recognition by the S1/S2 domain of SARS-CoV2 a structural analysis was performed by PyMOL. This is an approach combined with the molecular docking output files to analyze the interactions between the ligand and its receptor. We evaluat**e** the complexes obtained by docking to monitor intermolecular hydrogen bonds, electrostatic, and hydrophobic interactions between SARS-CoV2 S protein and TMPRSS2 missense variants compared to the wild type.

### Molecular Dynamics

MD simulations were performed using NAMD2 as a molecular dynamic program and VMD as a visualization program to understand the dynamic changes in the conformations of the wild type and missense variants-domain S1/S2 spike protein complexes in conditions close to those *in vivo*. MD simulations were carried out in water for 120 ns at constant temperature of 300 K, using the Langevin dynamics with a damping constant of 1 ps^−1^. The conformational changes observed during the simulation time frame are discussed below. Furthermore, VMD was used to determine the stability and mechanistic aspects of the wild type and mutant complexes by comparing their corresponding backbone root-mean-square deviation (RMSD), root-mean-square fluctuations (RMSF) and radius of gyration (Rg).

### MM−PBSA binding free energy

MD trajectories were used to compute the binding free energy of TMPRSS2 and missense variants to spike protein, using the molecular mechanics Poisson-Boltzmann solvent accessible surface area (MM-PBSA) method. This method is one of the most used approaches to estimate the free energy of binding of small ligands to biological macromolecules, it has been increasingly used in the study of biomolecular interactions. The total binding free energy (Δ*G*_*binding*_) can be calculated using Equation 1:

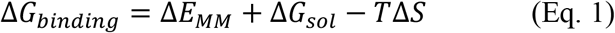

The MM/PBSA was used a fast and accurate method to predict the changes of binding free energy of the protein-protein complex caused by single point mutation. The effect of the polar and non-polar part of the solvent on the free energy was determined using the Poisson-Boltzmann equation and calculating the surface area. For our calculation, the outer dielectric constant was set to 80.0, the inner dielectric constant was set to 1.0, and the inverse of the grid spacing of 0.5 Å was used, while for the calculation of surface area, the surface tension value was set to 0.00542 with a surface offset of 0.92. And finally, the binding energy was summed and averaged over 10 snapshots.

## RESULTS AND DISCUSSION

### Polymorphism and molecular model of human TMPRSS2

To understand the role of TMPRSS2 variants in the infection by SARS-CoV2 virus, we searched on the Genome Aggregation Database (gnomAD), Ensembl, the 1000 Genomes Project and NHLBI databases to identify all SNPs in the *TMPRSS2* gene, causing amino acid changes at the protein level. Within the scoring ranges of the prediction tools, we identified a total of 692 missense and damaging SNPs with a relatively high allele frequency between 0.01 and 0.5.

Human TMPRSS2 is an 492 amino acid long protein with a transmembrane domain [TM] (84-106) and three functional domains: an N-terminal LDL-receptor class A domain [LDLRA] (133-148), followed by the cystein rich domain of the scavanger receptor [SRCR] (153-246) and finally at C-terminal peptidase S1 domain spanning from 256 to 492 amino acid which contains the protease active site residues: H296, D345 and S441. The catalytic domain (C-terminal peptidase S1 domain spanning from 256 to 492) of the crystal structure of human TMPRSS2, which is the only domain explored in our study, resolved using I-TASSER, has been aligned with the one recently deposited in the Protein Data Bank (code PDB: 7MEQ).

Figure 1 shows the alignment result of the two structures which shows a strong similarity between the two domains with an RMSD value equal to 0.705 Å, this proves that our model is well reliable.

**Figure1:**
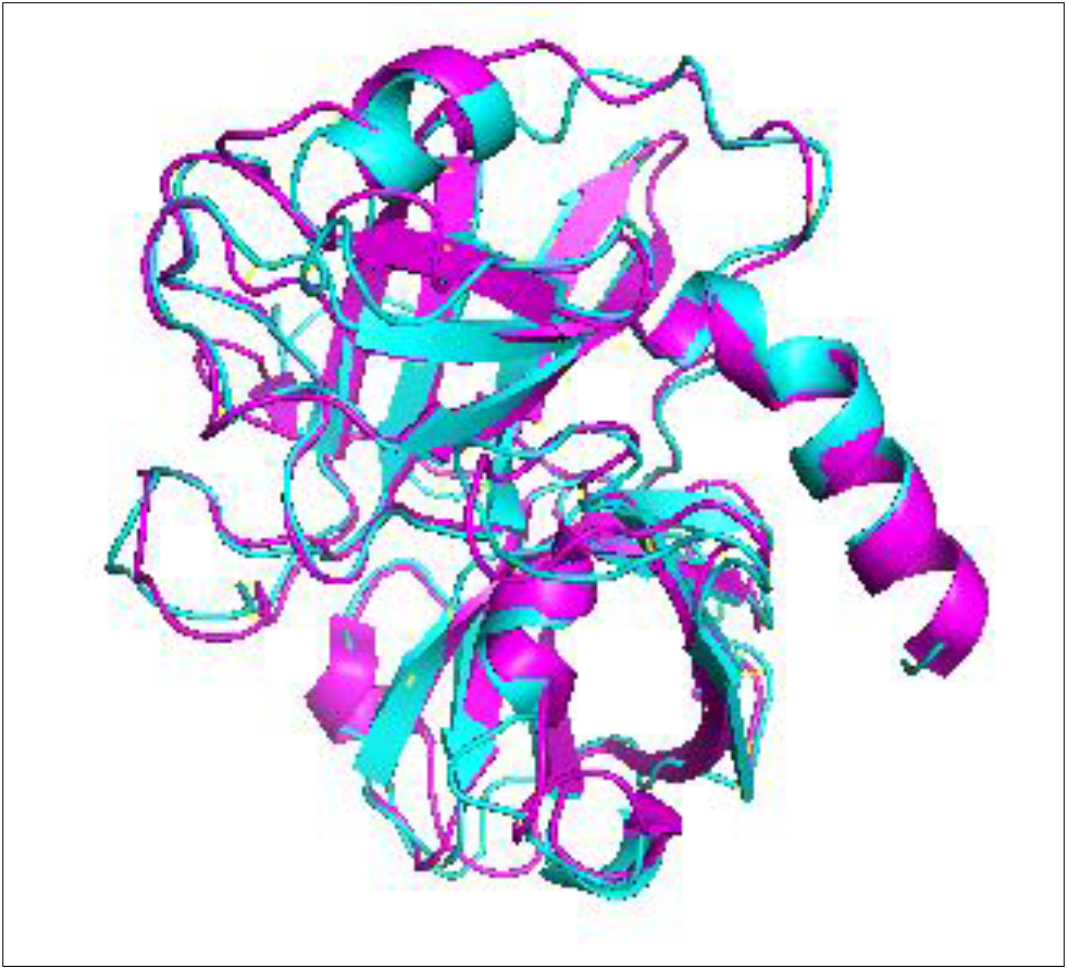
Comparison of the catalytic domain (256-492) of TMPRSS2 modelled by I-TASSER (shown in Cyan) with the model recently deposited in PDB (code PDB: 7MEQ shown in Magenta).

### TMPRSS2 binding interfaces

After obtaining the 3D structure of the TMPRSS2 protein, we mapped the catalytic triad of TMPRSS2, where residues H296, D345 and S441 were found to be highly conserved with other protease. Afterward, we predicted seven TMPRSS2 binding loops that may be involved in the interaction of the S protein with the TMPRSS2 protein, this identification is based on their position relative the TMPRSS2 protein catalytic triad (Figure 2.A). The different loops are as follow: Loop1: [Leucine273-Valine283], Loop2: [Valine298-Asparagine303], Loop3: [Tyrosine337-Asparagine343], Loop4: [Tryptophane384-Glutamic acid 395], Loop 5: [Serine412-Tyrosine416], Loop6: [**Aspartic acid 435-Aspartic acid 440]**, Loop 7: [Tryptophane461-Arginine470].

**Figure 2.**
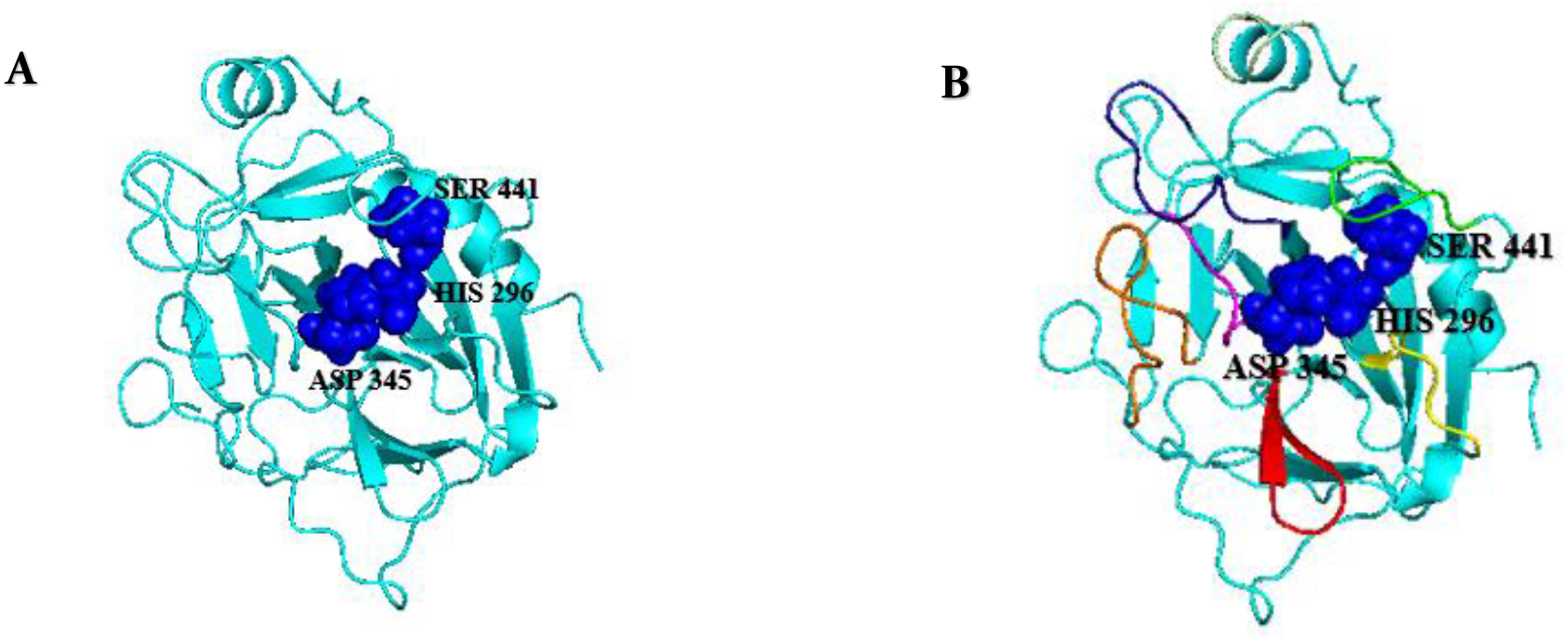
A) Secondary structure of the catalytic domain of TMPRSS2 protein and the TMPRSS2 catalytic triad (H296; D345; S441) are shown in blue spheres and B) Representation of TMPRSS2 protein binding loops.

**Figure 3.**
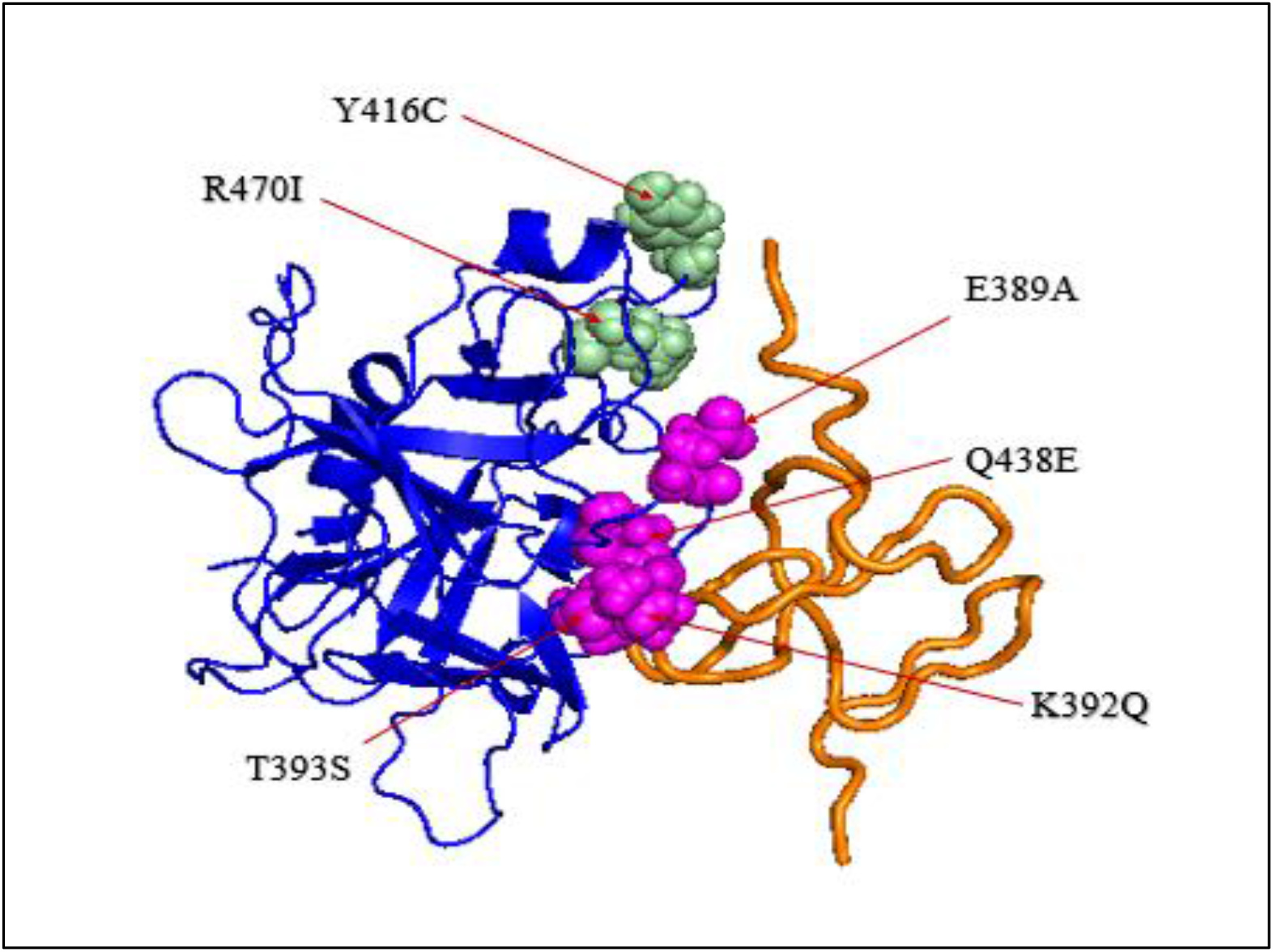
Secondary structure of the catalytic domain of TMPRSS2 protein (shown in blue) and the domain S1/S2 of the SARS-CoV2 (shown in orange). The mutated amino acids that “directly increasing” are colored in magenta and those “directly decreasing” are presented as green spheres.

### Polymorphisms of the TMPRSS2 gene related to the protein binding region to the viral particle and geographical distribution of this TMPRSS2 SNPs

The simplest way that missense variation could impact SARS-CoV-2 infection would be by altering the TMPRSS2-S interface. TMPRSS2 missense variants located at residues that bind the S protein are most likely to have such effects. In this study, among the 692 natural TMPRSS2 coding variants identified from the different databases, we found that 17 ponctual mutations at positions that have shown to be important for the binding of TMPRSS2 with the viral spike protein. Furthermore, GnomAD-Exomes database was used to gain information about frequencies of the examined TMPRSS2 SNPs worldwide. The population and the frequencies of each TMPRSS2 missense variants are plotted individually in the Table 1.

**Table 1:**
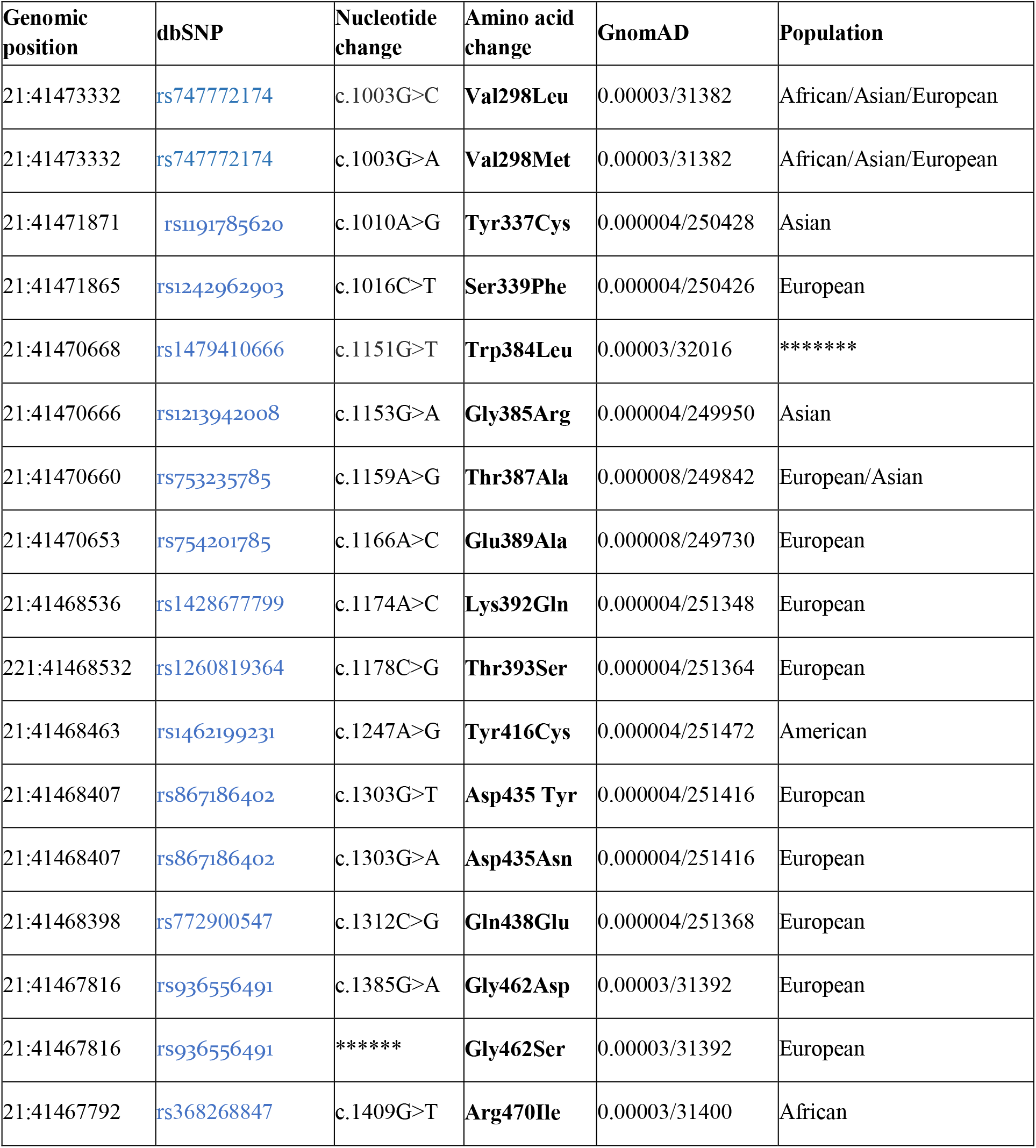
TMPRSS2 SNPs selected with their allele frequencies in each population

### 3D structures of the selected variants modelled by SWISS-MODEL

Structurally, all TMPRSS2 variants bear the characteristic domains of TMPRSS2 wild type. The overall protein architecture of TMPRSS2 allelic variants is largely conserved. The 3D structure of the 17 TMPRSS2 SNPs presents a significant similarity with the 3D structure of the wild type and the Ramachandran analysis of the different analogues proves that all the amino acids of the models are found in the favorable regions.

### Molecular docking of SARS-CoV2 S protein and TMPRSS2 wild type or missense variants

To analyze and quantify the binding affinities and interactions of different models of the 17 TMPRSS2 SNPs with the S1/S2 domain of the SARS-CoV2 S glycoprotein, we performed 18 docking simulations using AutoDock Vina software of theTMPRSS2 protein and the 17 SNPs with the viral glycoprotein and with the wild type protein and obtained the corresponding protein-ligand complexes. The binding energies are determined and reported in kcal/mol units in Table 2, which represents the binding energy of the best model identified with the docking software. Our docking simulations showed that TMPRSS2 and missense variants have high binding affinities with the domain S1/S2 of spike protein. Overall, the variants have a slightly higher interaction energies (0.3 to 1.1 kcal/mol) in respect to the wild type protein.

**Table 2:** Molecular docking results of the TMPRSS2 wild type and the corresponding 17 SNPs with the viral S protein.

| Protein position | Reference residue | Altered residue | Energy (kcal/mol) |
| --- | --- | --- | --- |
| TMPRSS2 | wild type | wild type | -13.8 |
| 438 | Q | E | -14.9 |
| 462 | G | S | -14.7 |
| 462 | G | D | -14.9 |
| 385 | G | R | -14.6 |
| 387 | T | A | -14.5 |
| 389 | E | A | -14.9 |
| 392 | K | Q | -14.9 |
| 393 | T | S | -14.5 |
| 416 | Y | C | -14.5 |
| 435 | D | Y | -14.7 |
| 435 | D | N | -14.7 |
| 470 | R | I | -14.6 |
| 298 | V | L | -14.3 |
| 298 | V | M | -14.1 |
| 337 | Y | C | -14.4 |
| 339 | S | F | -14.9 |
| 384 | W | L | -14.5 |

### Missense variants impact on binding of TMPRSS2 receptor to the viral S protein

Firstly, the interaction of wild type TMPRSS2 and the selected missense variants with the 3D structure of spike monomer protein were simulated using Autodock Vina. Additionally, both hydrogen, electrostatic bonds and hydrophobic bonds within different TMPRSS2-spike protein complexes were assessed using PyMOL. We classified the polymorphisms into two categories. The first one includes mutations that directly increase the interaction affinity within the complex, these variants increase the number of electrostatic interactions or decrease the distance of interaction between the receptor and ligand residues. We term these mutations as “directly increasing”. The second one includes mutations that directly decrease the interaction affinity of the complex by decreasing the number of electrostatic interactions or increasing the distance of interaction between the residues of receptor and its ligand, we term these mutations as “directly decreasing”. All these mutants were found to affect the residues in the TMPRSS2/S protein binding interface and in direct contact with the virus domain S1/S2 residues.

Therefore, this structure analysis allowed us to identify four missense variants E389A, K392Q, T393S and Q438E “directly increasing” the interaction affinity and 2 missense variants R470I and Y416C “directly decreasing” it.

### MD study

The complexes of the binding site (255-492) of TMPRSS2 and missense variants with the spike protein were subject of MD simulation studies over a period of 120 ns to understand their stability and study the structural consequences of these substitutions. To determine the stability and mechanistic aspects of the wild type and mutant complexes, hydrogen bond interactions, RMSD, RMSF, Rg and their binding profiles were analyzed and discussed below.

### Analysis of RMSD

The RMSD is usually used to measure the protein drift from a reference structure, to study the residue behavior of the protein during the simulations and to describe the dynamic stability of systems as it measures the global fluctuations of proteins or complexes. It reflected the mobility of an atom during the MD simulation trajectory. As a result, a higher residue RMSD value suggests higher mobility; inversely, a lower residue RMSD value suggests lower mobility. Therefore, the RMSD analysis was carried out for the MD simulations of each system to determine the change in the overall stability of the protein after mutation, more specifically to understand the effect of TMPRSS2 missense variants on the stability of complexes. In addition, we compared the RMSD of the wild type and mutants TMPRSS2/S protein complexes to the free forms of receptor (Figure S1) and the wild type complex to mutants TMPRSS2/S protein complexes (Figure S2) during the 120 ns of MD simulations.

### Analysis of RMSF

Protein RMSF was plotted to characterize local changes along the protein chain and to determine the movement of certain amino acid residues around their mean position to assess the flexibility of the dynamic nature of the residues during amino acid substitution, as a result, peaks indicate areas of the protein that fluctuated the most during the molecular dynamic simulation. It is well established that the flexibility determines the binding like it may not only affect the binding interface between two interacting partners but also an essential contribution to the entropy penalty during binding (Tuffery et Derreumaux, 2012).

RMSF of all the residues of the binding site (255-492) of the protein in each complex (TMPRSS2(WT), E389A, K392Q, T393S, Y416C, Q438E and R470I) in comparison to the RMSF of free forms of receptors have been calculated and plotted in (Figure S3) to understand the role of the amino acid substitution in the interaction with the S1/S2 domain of Spike protein of SARS-CoV-2.

### Analysis of gyration radius

Radius of gyration measures the compactness and the dimension of protein-protein complex during MD simulations that shows the stability of protein folding. We performed Rg analysis to observe the conformational alterations and dynamic stability of the TMPRSS2(WT)-S1/S2 domain of spike protein and their corresponding mutant complexes. Data are displayed in Figure S4.

### Dynamics of hydrogen bonds

Analysis of the hydrogen bonds (HB) during ligand binding is another important factor that influences protein stability, it has a significant role to strengthen protein-protein interactions. To elucidate how the mutations affect the TMPRSS2 and viral protein interaction at molecular level, the dynamics of hydrogen bonds of each system is displayed in Figure S5, followed by the evolution of the HB number of each complex and free forms of the receptor during 120 ns of MD simulations (Figure S6), finishing with a comparison of the dynamics of hydrogen bonds of each mutant system with the WT complex (Figure S7).

### Analysis of the binding free energy

Furthermore, to understand and quantify the strength of the interactions between a ligand and protein, the binding energies over 10 snapshots is reported as the final ΔG_binding_. The binding free energy calculation of protein–ligand complexes is necessary for research into virus–host interactions. Based on the MD simulation trajectories, the binding free energies of TMPRSS2 (WT) and selected missense variants to S protein were calculated using the MM-PBSA method (Ben shalom et al., 2017) that may ignore the change in structure of the ligand and the receptor upon ligand binding (Genheden et Ryde, 2015), which may be important factors for the affinity. As a result, the negative energy of the binding complex shows the strength of the protein–ligand interaction.

Table 3 shows a comparison between the free energies of the TMPRSS2 proteins for SARS-CoV2. Analysis of the results showed that the binding free energy of -39.1 kcal/mol corresponds to the binding energy of the TMPRSS2 WT which is the highest value compared to other complexes. In the other hand, the lowest binding free energies of Q438E, Y416C, T393S, R470I, E389A and K392Q to S1/S2 domain of S protein are –42.7, –54.0, –61.0, –61.2, –75.1 and –82.8 kcal/mol, respectively. All the variants have a higher binding energy than the native does, this result suggests that the missense variants have stronger binding affinity that can be explained by the strong affinity of these variants towards the S protein compared to the native TMPRSS2.

**Table 3:** MM/PBSA binding free energies (kcal/mol) of wild-type and mutant complexes

| Models | Complex | Receptor | Ligand | $\Delta G_{binding}$ ( kcal/mol) |
| --- | --- | --- | --- | --- |
| Native | -7573.2 | -5624.8 | -1909.2 | -39.1 |
| Q438E | -7601.7 | -5667.4 | -1891.6 | -42.7 |
| Y416C | -7553.3 | -5610.1 | -1889.2 | -54.0 |
| T393S | -7525.9 | -5558.9 | -1906.0 | -61.0 |
| R470I | -7306.2 | -5343.4 | -1901.5 | -61.2 |
| E389A | -7546.5 | -5563.6 | -1907.8 | -75.1 |
| K392Q | -7529.1 | -5591.1 | -1855.2 | -82.8 |

## DISCUSSION

Based on recent reports, TMPRSS2 is essential for SARS CoV2 to enter cells, it is one of the main cell surface proteases involved in the process of S protein priming. However, to the virus can enter to the cells, a first cleavage of the viral spike protein at the S1/S2 site that is very important for the activation of virus, followed by second cleavage at the S2’ site, which allows viral fusion with the cell membrane and internalization.

The gene encoding TMPRSS2 has a high level of genetic variability. In this context (Yuan et al., 2020), (Ravikanth et al., 2020), (Paniri A et al., 2020), (Senapati S et al., 2020) suggested that TMPRSS2 DNA polymorphisms were likely to be associated with susceptibility to COVID-19 and would contribute to differences in SARS-CoV2 infection. In the other hand, recent studies showed that *ACE2* genetic variation is very rare in the population (Stawiski et al.) (MacGowan and Barton), thus making it a candidate to explain the inter-individual variability to SARS-CoV2 infection. At present, no disease-association for *TMPRSS2* variants is known. Therefore, we focused on *TMPRSS2*, which together with *ACE2* plays an important role in SARS-CoV2 infection. Our *in-silico* analysis of human *TMPRSS2* variants was carried out to verify the hypothesis that the COVID-19 susceptibility is also influenced by genetic variability of gene coding for TMPRSS2 protein involved in the entry of SARS-CoV-2 into target cells and that certain populations may be more affected by SARS-CoV2, depending on the frequency of TMPRSS2 variants. Hence, for further understanding of the susceptibility of individuals of different populations to SARS-CoV2 and their risk of infection, we analyzed the 1000Genomes, Ensembl, NHLBI, genomAD databases dedicated to mutations to extract the missense variants of the TMPRSS2 protein. In total, 642 missense variants were obtained. After modelling the TMPRSS2 3D structure using I-TASSER, we have compared the catalytic domain of our structure to the one recently deposited in PDB (code PDB: 7MEQ), which shows a strong similarity with an RMSD value equal to 0.705 Å, then we predicted the binding site with the S1/S2 domain of the S protein. In a further step, we focused only on the missense variants whose spatial position is at the level of the binding site in order to identify those that are able to modify the interaction affinity in a direct way with the S protein.

In the present study, we performed an *in-silico* analysis of the SNP variants localized at the binding loops. Those SNPs can be directly involved in the alteration of interaction affinity based on molecular docking to obtain the complexes of TMPRSS2. We also selected missense variants with the viral protein to predict the interaction affinity between the two partners, followed by a structure function analysis to identify the key bonds of interaction. In a last step, we carried out a MD study for the wild-type complex and variants that have the potential to alter the interaction affinity between TMPRSS2 and the S protein with aiming to validate the previous results and identify missense variants of TMPRSS2 that can alter the interaction affinity with the viral protein relative to the native complex. To determine the stability and mechanistic aspects of the wild type and mutant complexes, HB interactions, RMSD, RMSF, Rg and their binding profiles were analyzed.

On the other hand, Senapati et al and Ravikanth et al suggested that variants in TMPRSS2 that are considered damaging by prediction tools may alter the structure of TMPRSS2 which may indirectly affect the interaction affinity with SARS-CoV2 spike protein through structural change (Ravikanth et al., 2020; Senapati et al., 2020). Add to that, Hussein et al performed an *in-silico* study and tested the effect of the frequent V160M mutation, which is localized at the serine protease domain and suggested that this mutation can indirectly modify the interaction affinity (Hussein et al.,2020).

The results reported in this study show a remarkable change in the interaction affinity of the missense variants with the spike protein compared to the native protein and suggest that these missense variants may be directly involved in the modification of interaction affinity between the human TMPRSS2 protein and SARS-CoV2. The binding free energy of the 6 SNP variants is higher than that of the native one but the two variants Y146C and R470I, which are considered by the structure function analysis as decreasing the binding affinity have a less stable RMSD compared to native, which can decrease the stability of the two complexes. The R470I is present only in the African population with an allelic frequency equal to 0.00003 and the Y416C is present only in the American population with an allelic frequency equal to 0. 000004. However, the 4 SNP variants E389A; K392Q; T393S and Q438E which are considered to increase the interaction affinity are present only in the European population with allelic frequencies equal to 0.000008;0.000004;0.000004;0.000004, respectively.

## CONCLUSION

The COVID-19 pandemic highlighted the functional role of the TMPRSS2 protein in the priming of the SARS-CoV2 spike protein and the internalization of the virus inside the host cell. TMPRSS2 is an essential component for viral infection, allowing the activation of the S protein by cleaving it to generate two distinct fragments. It may therefore be a potential target for the development of therapeutic and preventive approaches. Several studies have shown the inter-individual variability to SARS-CoV2 infection highlighting the involvement of demographic, environmental and genetic factors. Natural genetic variations in the TMPRSS2 gene can modulate the affinity of the interaction of the TMPRSS2 receptor and the SARS-CoV-2/S protein and lead to a difference in the susceptibility of the virus response. Our data suggests that certain populations might be more affected by SARS-CoV2, depending on the frequency of the respective variants. Overall, the mutations identified in TMPRSS2 human protein binding domain to SARS-CoV2 had led to structural changes with modification of the interaction affinity between the TMPRSS2 receptor and spike protein. The RMSD, RMSF, Rg and HB number of the 120 ns simulation run confirms the modification of stability caused by TMPRSS2 missense variants in comparison to the wild type one. The energy calculations reiterate the binding efficiency of missense variants in comparison to the wild type. Finally, our results can potentially guide future attempts, to design an inhibitor containing TMPRSS2 missense variants that are capable of increasing interaction affinity with spike protein to disrupt the interaction between the TMPRSS2 human protein and SARS-CoV2.

## ACKNOWLEDGEMENTS

For computer time, this research (ref. **K1495**) used the resources of the Supercomputing Laboratory at King Abdullah University of Science & Technology (KAUST) in Thuwal, Saudi Arabia. The IAU DSR project ID is 2021-113-Sci, The KAUST project ID is K1495 and The Tunisian federated research project ID is PRFCOV19-D.2P2.

